# The Dual Roles of the CXCL10-CXCR3 Axis and Its Therapeutic Potential in Osteosarcoma

**DOI:** 10.1101/2024.06.04.597467

**Authors:** Benjamin B. Gyau, Junyan Wang, Xiang Chen, Margaret Clement, Zoe D. Man, Angela Major, Mathew Weiser, Jun Xu, John Hicks, Tsz-Kwong Man

**Author notes:** Corresponding author: Tsz-Kwong Man, Associate Professor, Baylor College of Medicine Mailing address: Suite 1200, 1102 Bates Ave, Houston, TX 77030.

## Abstract

The CXCL10-CXCR3 axis is recognized for its dual role in tumor biology, promoting tumor growth and metastasis via autocrine signaling while also eliciting anti-tumor responses through paracrine signaling. However, its specific functions in osteosarcoma (OS), the most prevalent malignant bone tumor in children, remain poorly understood. Our previous research has demonstrated that elevated circulating CXCL10 levels correlate with poor prognosis in OS patients. Analysis of the TARGET OS RNAseq dataset revealed that high expression levels of CXCL10 or its receptor CXCR3 are associated with improved prognosis. Given the known role of CXCL10 in recruiting CXCR3+ immune cells to combat cancer, we further analyzed single-cell RNAseq data and found that CXCR3 is predominantly expressed in CD3+ T cell populations. These findings suggest that CXCL10 may also play a protective role in OS by recruiting anti-tumor immune cells. To elucidate the causal role of the CXCL10-CXCR3 axis in OS, we conducted *in vitro* phenotypic assays on three OS cell lines with and without CXCL10. The chemokine was found to enhance tumor cell migration and AKT phosphorylation. Utilizing a CRISPR-mediated CXCR3 deletion mutant, we demonstrated that the absence of CXCR3 significantly inhibited OS tumor growth and pulmonary metastasis in an orthotopic xenograft mouse model. Transfection with the CXCR3A isoform, but not the CXCR3B isoform, restored the migratory phenotype of the CXCR3 deletion mutant to levels comparable to the parental cell line. Additionally, pharmacological inhibition of CXCR3 with AMG487 markedly reduced OS cell migration *in vitro* and metastasis development in the orthotopic xenograft mouse model. Our research highlights the complex interplay of the CXCL10-CXCR3 axis in both tumor and immune cells. We propose a working model for the roles of the CXCL10-CXCR3 axis in OS, suggesting that targeting CXCR3 may be an effective strategy to inhibit OS metastasis, particularly in immune-cold OS subtypes.

## INTRODUCTION

Osteosarcoma (OS) is a rare yet devastating malignant bone tumor affecting children primarily in the second decade of life. Despite significant medical advancements, survival rates for OS have seen little improvement over the past four decades. The primary treatment modalities for children with OS are surgery and conventional chemotherapy.^1–3^ However, complete eradication of cancerous cells through surgery is seldom achieved, providing opportunities for tumor metastasis. Metastasis represents the leading cause of mortality in OS patients, with survival rates for metastatic OS currently hovering below 25%.^4,5^ Conventional chemotherapeutic agents like cisplatin, doxorubicin, and high-dose methotrexate, while commonly used, often come with severe side effects that significantly impact the quality of life for affected children. Moreover, since effective therapies for metastatic OS remain elusive, all patients are subjected to the same chemotherapy regimen regardless of their individual clinical risks of metastasis.^6,7^ There exists a pressing need for more efficacious treatment approaches with reduced side effects to combat OS. A deeper understanding of the key drivers of metastasis holds the potential to unveil novel therapeutic strategies that can be further evaluated in clinical settings.

The C-X-C motif family of chemokines and their corresponding receptors are pivotal in tumor development and immune modulation.^8–10^ For instance, in pancreatic adenocarcinoma, NF-kB-dependent expression of CXCL5 fosters tumor progression by recruiting CXCR2+ myeloid-derived suppressor cells (MDSCs).^11^ Likewise, in breast and prostate cancers, the CXCL12-CXCR4 and CXCL13-CXCR5 signaling axes are crucial for distant-organ metastasis.^12^ Notably, CXCR3 is prominently expressed by anti-tumor immune cells, including activated CD8+ T, NK, and B cells.^13–15^ Although involved in inflammation and immune cell trafficking, the CXCL10-CXCR3 axis has also been implicated in the development and advancement of various malignancies, including OS.^16–20^ In breast cancer, high CXCR3 expression correlates with a poorer prognosis,^21,22^ and antagonizing CXCR3 signaling in a murine model of metastatic breast cancer notably inhibits lung metastasis.^19^ Similarly, administration of the CXCR3 antagonist AMG487 significantly reduces metastatic colonization in lungs using experimental metastasis models of OS, but its effect on the metastatic cascade originating from the primary tumor is still elusive.^17^

Recently, we identified CXCL10 as a prognostic biomarker for OS and demonstrated its association with metastasis and poorer outcomes in OS patients.^23^ Building upon this, our current study extends previous research on CXCR3 in OS by investigating the roles of both CXCL10 and CXCR3 in OS metastasis. Additionally, we assessed the targetability of CXCR3 using a CRISPR-mediated deletion mutant and the CXCR3 antagonist AMG487 in orthotopic xenograft mouse models, which faithfully recapitulate the full metastatic cascade. Our findings underscore the critical role of CXCR3 signaling in OS metastasis development and highlight CXCR3 inhibition as a promising therapeutic strategy for controlling lung metastasis.

## MATERIALS AND METHODS

### Mice and human cell lines

Four-to-six-week-old NOD. CB17-Prkdc SCID/J mice were procured from Jackson Laboratory and housed in a pathogen-free facility at Baylor College of Medicine, Houston, TX. Mice were fed on a standard Clea Rodent CL-2 diet throughout the experiment. 143B, LM7, MG63.3 OS cell lines used in this study were initially purchased from the American Type Culture Collection and stored in liquid nitrogen until needed. Cells were maintained in DMEM supplemented with 10% FBS. The luciferase-labeled 143B cell line utilized in this study has previously been reported.^24^

### CRISPR-mediated CXCR3 deletion mutant (CXCR3 KO) generation

143B-Luc cells were grown in DMEM with 10% FBS and 1% Penicillin/Streptomycin. Cells were trypsinized and passaged 1 day before electroporation. On the day of electroporation, 30 pmol of 2 sgRNAs (sgRNA1: agugcuaaaugacgccgagg, sgRNA2: cuacgcaggagcccuccugc, from Synthego) were mixed with 10 pmol Cas9 protein (Integrated DNA Technologies) in 2 µL of Resuspension Buffer R (Neon Electroporation Kit, Cat# MPK1096, ThermoFisher) and incubated at room temperature for 20 minutes to form the CRISPR RNP complex *in vitro*. 100,000 cells were electroporated with the assembled RNP at 1200v for 30 minutes and 1 pulse using the NEON electroporator (ThermoFisher). After electroporation, cells were plated in 1 mL of media and allowed to recover for 48 hours. Cells were then trypsinized and diluted to ∼50 cells per 10 mL of media, and plated at 100 µL per well in 96-well plates. When the cells grew back from single cells after 1 to 2 weeks of incubation, they were moved to 24-well plates where they were further monitored for growth. Once the cells were confluent enough (>50%), they were collected and screened for deletion by genomic PCR.

### Polymerase chain reaction

Genomic DNA was extracted using QuickExtract DNA extraction solution (Biosearch Technologies) and PCR was performed with OneTaq Standard 5X buffer and OneTaq Polymerase (New England Biolabs) using a 96-well Thermal Cycler (ThermoFisher). PCR primers used were fwd-5’atgcgagagaagcagccttt and rev-5’gaagtctgggagggcgaaaa (Integrated DNA Technologies). PCR amplifications consisted of an initial denaturation at 94°C for 2 minutes, followed by 32 cycles of 94°C for 30 seconds, 53°C for 30 seconds, and 68°C for 1 minute. The samples with expected deletion (∼400bp) in the PCR screen were then treated with Shrimp Alkaline Phosphatase and Exonuclease I (New England Biolabs) and sent out for Sanger sequencing (Eurofins Genomics).

### CXCR3A and CXCR3B expression construct transfection

CXCR3 KO cells were seeded overnight at a density of 2.5 × 10^5^ cells per well in a 6-well plate. Cells were then transfected with 3 μg of CXCR3A plasmids (GenScript, OHU18425C) or CXCR3B expression construct (GenScript, OHU16413). Each plasmid was mixed with 9 μL of Lipofectamine2000 (Invitrogen, UA) and incubated for 20 minutes at room temperature before being transferred into the growth medium. After 48 hours of transfection, stable clones were selected and isolated in DMEM with 10% FBS and 1 mg/ml G418. Successful transfections were confirmed by Western blotting and PCR.

### Immunoblotting

OS cell lines were grown in DMEM supplemented with 10% FBS and 1% Penicillin/Streptomycin. At 80% confluence, cells were harvested and lysed in ice-cold RIPA buffer containing 100X protease and phosphatase inhibitors (Cell Signaling). Protein concentrations were determined using BSA protein assay (Bio-Rad). SDS-PAGE electrophoresis and Western blotting were conducted as previously described.^25^ The primary antibodies used were: CXCR3 (1:1000, Invitrogen, PA5-28741), Vinculin (1:5000, Sigma, V9131), P-AKT^S473^ (1:1000, Cell Signaling, 4060), GAPDH (1:10000, Santa Cruz, sc-32233), CXCL10 (1:1000, Cell Signaling, 14969). Secondary antibodies used were anti-rabbit and anti-mouse IgG HRP-linked antibodies (1:5000, Cell Signaling, 7074 and 7076 respectively). Images were developed using the MINI-MED 90 X-Ray Film Processor (AFP Manufacturing Corp). Protein bands were visualized using chemiluminescent substrates (Thermo Fisher) on autoradiography films with a pour developer (Merry X-Ray Corp).

### Cell migration assay

The QCM 24-well cell migration assay kit (ECM508, Millipore) was used to assess the CXCL10-mediated chemotaxis of OS cells according to the manufacturer’s instructions. Briefly, cells were starved overnight and assayed in 0.1% FBS-supplemented DMEM (FBS-depleted medium) except indicated otherwise. A total of 300 µL of 5 x 10^4^ cells was seeded in the insert well while 500 µL of media with or without various concentrations of CXCL4, CXCL9, CXCL10, and CXCL11 were added to the lower chamber. 10% FBS-supplemented DMEM in the lower chamber served as the positive control. In the AMG487 experiment, cells were pre-treated with 1 µM AMG487 for 1 hour before incubation with or without 100 ng/mL CXCL10. After 4 hours of incubation at 37°C in a 5% CO_2_ incubator, migrated cells were stained with crystal violet dye Cell Stain, imaged at 4X magnification (EVOS FL Microscope, Invitrogen), and quantified with ImageJ (v1.53e; Java 1.8.0_172). Results from five random fields per well were analyzed and averaged. Triplicate independent experiments were performed. The chemotactic index, CI, was calculated as the ratio of the migrated cells in the CXCL10-stimulated wells to the unstimulated wells.

### Cell proliferation assay

A total of 10^4^ OS cells starved overnight in an FBS-depleted medium were seeded and treated in 0.1% FBS-supplemented DMEM (depleted medium) in 96-well plates. Cell viability was assessed using a CCK-8 kit (Dojindo Laboratories). 10 µL CCK-8 solution was added to each well, and incubated for 1 hour at 37°C. Absorbance was read at 450 nm with the MiniMax 300 imaging cytometer (Molecular Devices).

### Mouse studies

10^6^ 143B-Luc cells were orthotopically injected into the tibia of eight-to-sixteen-week-old NOD/SCID mice. 9 (4 males and 5 females) and 8 (4 males and 4 females) mice were injected with 143B CXCR3 parental (WT) and CXCR3 deletion (KO) cells respectively. When tumors were palpable, approximately 2 weeks after injection, mice were imaged once weekly with IVIS to monitor tumor growth. 5 weeks post-surgery, mice were euthanized for histopathological examination of lung tissues to quantify pulmonary metastases. A similar experimental design was adopted for the rescue experiments by injecting KO cells stably transfected with CXCR3A (7 mice; 4 males and 3 females) and CXCR3B (6 mice; 3 males and 3 females) isoforms. For the AMG487 experiment, tumor-bearing mice were randomly divided into two groups two weeks after intratibial injection. The treatment group (6 mice; 3 males and 3 females) received a twice daily subcutaneous injection of 100 µL of 5 mg/kg mouse weight of AMG487 (Tocris, Cat# 448710) for 2 weeks, and the vehicle group (8 mice; 4 males and 4 females) had the same injection volume, frequency, and duration of 20% hydroxypropyl-ß-cyclodextrin (the drug vehicle). Mice were imaged once weekly with IVIS to monitor tumor growth and were euthanized after the treatment. Lung tissues were then harvested for histological analysis to analyze pulmonary metastases.

### Statistical and bioinformatic analyses

For the chemotactic index analysis, the statistical significance was computed using one-tailed, one-sample t-tests with a reference value of 1. The results from CCK-8 assays and mouse studies are presented as mean ± SEM. To assess differences in pulmonary metastases, two-tailed Mann-Whitney tests were employed. Additionally, two-way analysis of variance (ANOVA) was used to evaluate the statistical significance of primary tumor growth in mouse studies and cell proliferation in CCK-8 assays. GraphPad Prism v10.0 facilitated the statistical analyses, and a significance level of P < 0.05 was considered.

The RNAseq data used in this study are in whole or part based upon data generated by the Therapeutically Applicable Research to Generate Effective Treatments (TARGET) initiative, phs000218, managed by the NCI. The data used for this analysis are available for download through the Genomic Data Common (GDC). Survival analysis of the TARGET RNAseq data was performed with Kaplan-Meier analysis implemented in BRB-ArrayTools v4.6.2 developed by Dr. Richard Simon and the BRB-ArrayTools Development Team. Single-cell RNAseq data was analyzed by Seurat v5.01.^26^

## RESULTS

### Elevated CXCR3 expression is linked to improved prognosis and increased T cell infiltration

Our previous finding has shown the link between high circulating levels of CXCL10 with poor prognosis and metastasis in OS, suggesting a pro-metastatic role for this chemokine.^23^ To elucidate the prognostic role of CXCL10 and its receptor CXCR3 in OS tumors, we conducted a survival analysis on publicly available RNA expression data from the Therapeutically Applicable Research to Generate Effective Treatments (TARGET) OS cohort. Surprisingly, our analysis revealed that high levels of CXCL10 or CXCR3 were associated with better survival (Figure 1A-B), suggesting a protective role for this axis, in contrast to previous observations from circulating levels. To further understand the discrepancy between systemic and tumor-specific effects, we examined the expression of CXCL10 and CXCR3 in a previously published single-cell RNAseq OS dataset.^27,28^ Our analysis showed that CXCL10 expression was minimal in OS tumors, whereas CXCR3 was predominantly expressed in the CD3+ T cell population (Figure 1C). These findings suggest that the elevated CXCR3 expression observed in OS cases may be attributed to heightened T cell infiltration, thereby indicating a better prognosis.

**Figure 1:**
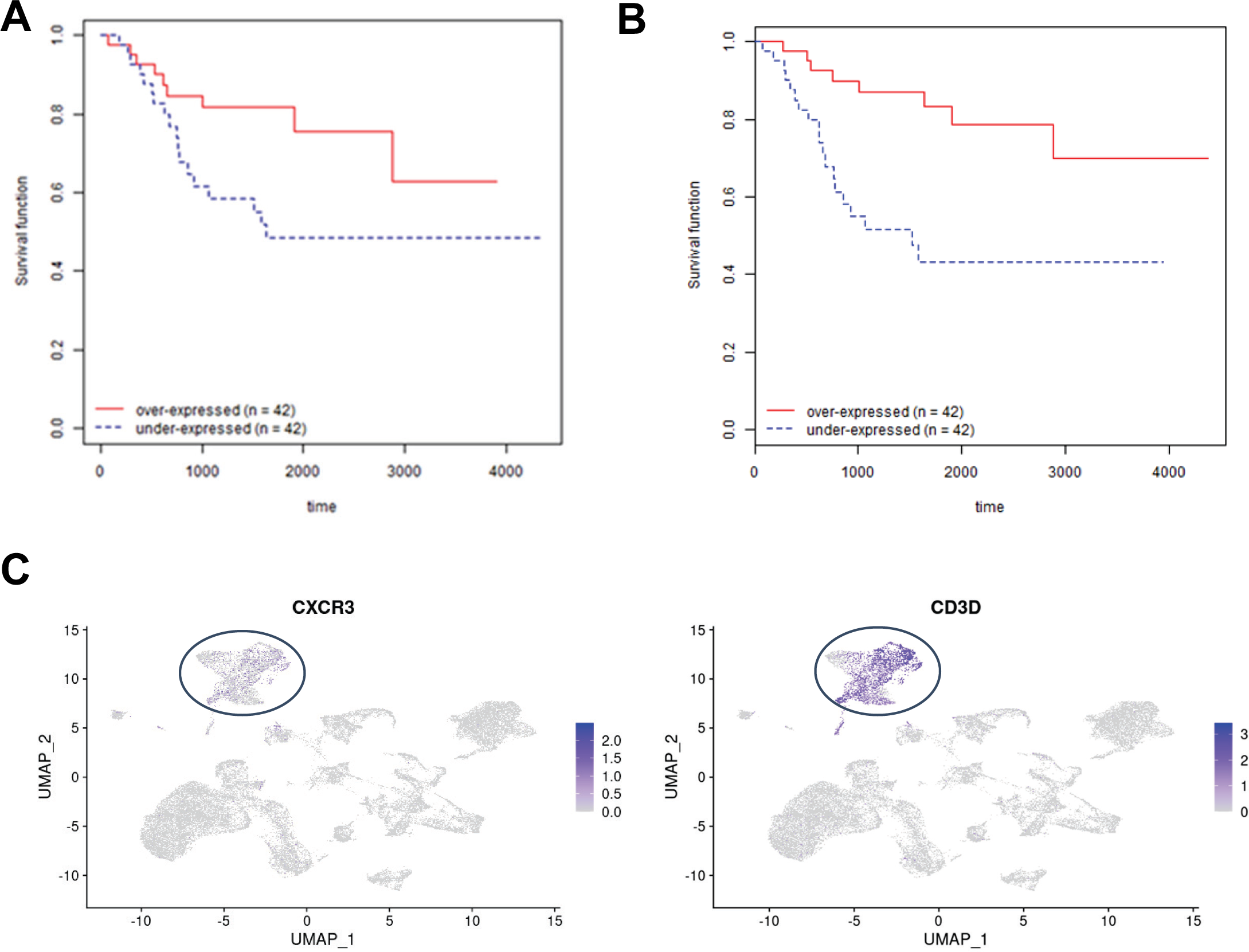
Kaplan-Meier and scRNAseq analyses of CXCR3 and CXCL10 expressions in OS. **A,** Overall survival analysis of RNA expression of CXCL10 and CXCR3 in the TARGET RNA expression dataset. Patients were classified into over-expressed (red solid) and under-expressed (blue dashed) groups based on the median expression of each transcript. Wald tests were used to determine significant differences between the two expression groups for each gene, with P < 0.05 considered significant. **B,** UMAPs of scRNAseq data from OS cases (GSE162454) showing CXCR3 is predominantly expressed in the CD3+ T cell population (circled).

### CXCL10 enhances OS cell migration and AKT phosphorylation

To elucidate whether the CXCL10-CXCR3 axis exhibits pro-metastatic or protective effects in OS, we investigated the migratory impact of exogenous CXCL10 on three commonly used metastatic OS cell lines—143B, LM7, and MG63.3—each with varying levels of CXCR3 expression (Figure 2A). In transwell migration assays, we observed a significant increase in OS cell migration with at least one of the four tested concentrations of CXCL10 compared to the control without CXCL10 as quantified by the chemotactic index (Figure 2B). In addition to CXCL10, CXCL4, CXCL9, and CXCL11 are also cognate CXCR3 ligands with differing receptor affinities and functional effects on target cells.^29–31^ Transwell migration results showed that CXCL4 and CXCL9 significantly enhanced cell migration in two of the three OS cell lines, while CXCL11 promoted migration in all three cell lines (Figure 2B). These findings suggest that the migratory effects of CXCR3 ligands are not exclusive to CXCL10. To elucidate the downstream signaling mechanism underlying CXCL10-induced migration, we conducted phospho-AKT immunoblotting in the three OS cell lines following incubation with various CXCL10 concentrations at two time points. AKT signaling has been shown to be downstream of CXCL10 signaling.^32–34^ Our results revealed that CXCL10 triggered an increase in AKT phosphorylation across all three cell lines compared to cells incubated with the depleted medium, particularly evident at the 120-minute time point (Figure 2C). This suggests that CXCL10-mediated OS chemotaxis is mediated via the AKT pathway.

**Figure 2:**
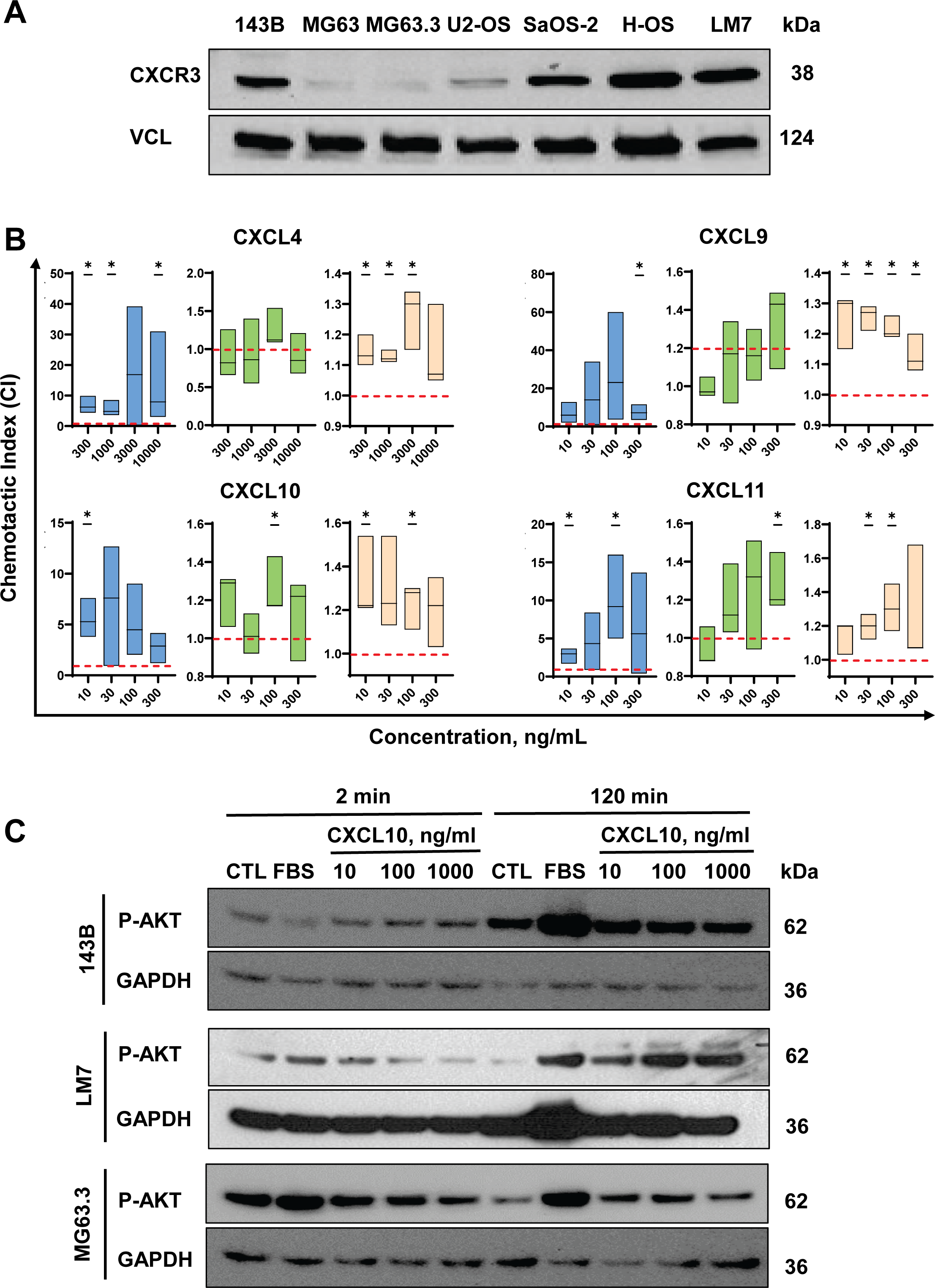
The effects of CXCL10 on OS cell migration and AKT phosphorylation. **A,** CXCR3 immunoblotting of seven OS cell lines maintained in DMEM supplemented with 10% FBS. Vinculin was used as a loading control. **B,** Results of transwell migration assays for three OS cell lines in response to CXCR3 ligands over 4-6 hours. The chemotactic index (CI) was calculated as the ratio of migrated cells in CXCL10-stimulated wells to those in unstimulated wells. **C,** Phospho-AKT immunoblotting of OS cell lines incubated with CXCL10. Three OS cell lines were treated with various concentrations of CXCL10 for two different time durations. GAPDH was used as a loading control. Cells incubated with 10% or 0.1% FBS served as positive and negative controls, respectively. Statistical significance was determined using one-tailed one-sample t-tests with a reference value of 1 (*P < 0.05; ns = not significant).

### Deletion of CXCR3 attenuates OS growth and metastasis in an orthotopic xenograft mouse model

To ascertain whether CXCL10-mediated AKT phosphorylation and OS cell migration rely on the CXCR3 receptor, we generated a CRISPR-mediated CXCR3 deletion mutant in 143B cells (CXCR3 KO). The sequence validation of the mutant is provided in Supplementary Figure S1. We further confirmed the gene deletion through CXCR3 immunoblotting (Figure 3A). Subsequent incubation of parental and KO cells with CXCL10 demonstrated a marked reduction in both AKT phosphorylation (Figure 3B) and OS migration (Figure 3C) in the KO mutants compared to parental 143B cells, underscoring the necessity of CXCR3 for CXCL10-mediated migration and AKT phosphorylation.

**Figure 3:**
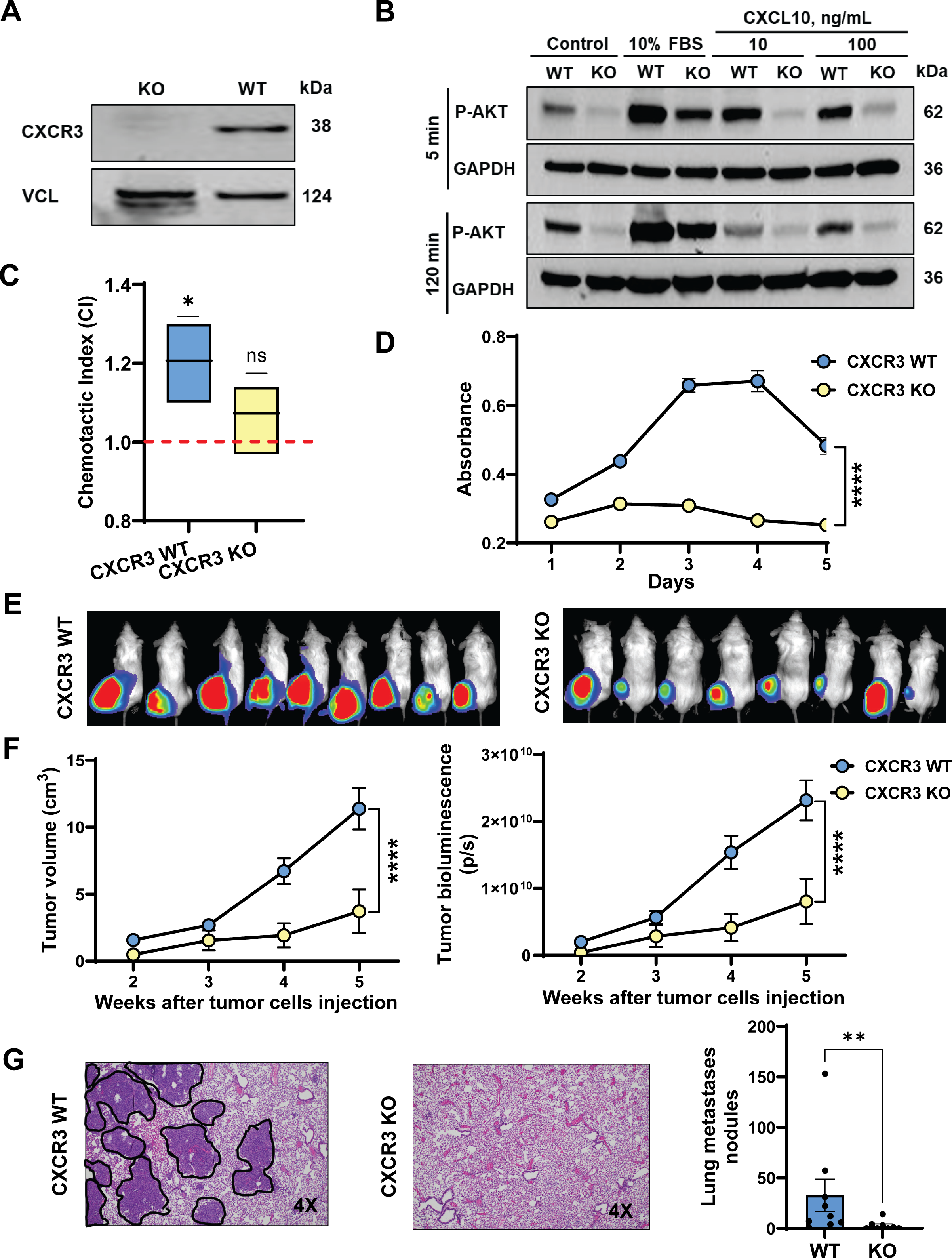
Orthotopic xenograft mouse model investigating the effects of CXCR3 deletion on the development of pulmonary metastasis in OS. **A,** CXCR3 immunoblotting of parental 143B (WT) and CRISPR-mediated CXCR3 deletion mutant (KO) cells. **B,** Phospho-AKT (P-AKT) immunoblotting of parental 143B and CXCR3 KO mutant cells exposed to CXCL10 for 5 minutes and 2 hours, with GAPDH as a loading control. Negative and positive control cells were treated with 0.1% FBS medium without CXCL10 and treated with 10% FBS medium, respectively. **C,** CXCL10-mediated transwell migration assays of parental 143B and CXCR3 KO mutant cells in 0.1% FBS DMEM. **D,** CCK8 proliferation assays of parental 143B and CXCR3 KO mutant cells in 0.1% FBS DMEM. **E,** IVIS images of orthotopic xenograft mouse models taken 5 weeks after intratibial injection with parental 143B (left panel) and CXCR3 KO mutant cells (right panel). **F,** Comparisons of tumor volumes, calculated as 0.5 x h x w² (left panel), and bioluminescence of primary tumors (right panel) for the mouse models. **G,** Representative images (left) and quantification (right) of metastatic nodules in H&E-stained resected lungs from the mouse models, with each dot denoting one mouse. Error bars and asterisks represent standard deviations and statistical significance (*P < 0.05; **P < 0.01; ***P < 0.001; ****P < 0.0001; ns = not significant).

While the distant-organ migratory propensity of most tumors is primarily determined by genetic factors, it is often preceded by tumor proliferation.^35,36^ To elucidate the role of CXCR3 in OS growth, we examined whether the absence of this chemokine receptor affected OS cell proliferation in a serum-depleted medium. Results from the CCK-8 proliferation assays of 143B and CXCR3 KO cells revealed a significant reduction in proliferation of the KO mutant compared to the parental cell line, suggesting that CXCR3 also contributes to OS cell proliferation (Figure 3D).

Building on our observation that the CXCL10-CXCR3 axis is vital for promoting OS migratory phenotypes *in vitro*, we conducted an *in vivo* analysis to assess the impact of CXCR3 deletion on tumor growth and metastasis development using orthotopic xenograft mouse models. Luciferase-labeled parental 143B or CXCR3 KO cells were orthotopically injected into the tibia of NOD/SCID mice. Tumor growth was monitored via IVIS over a 5-week period, and the extent of pulmonary metastasis was assessed through H&E staining of resected mouse lungs post-euthanization. The absence of tumoral CXCR3 significantly reduced both tumor volume and luciferase activity (Figure 3E-F), as well as the number and size of pulmonary metastatic nodules compared to mice injected with parental 143B cells (Figure 3G). Notably, there was no positive correlation between primary tumor size or activity and the rate of lung metastasis, suggesting that the distant-organ spreading of OS cells and primary tumor proliferation are distinct processes (Supplementary Figure S2).

### The CXCR3 deletion phenotypes are recused by the CXCR3A isoform

Among the two major transcript isoforms described in CXCR3, CXCR3A promotes tumor cell migration, while CXCR3B impedes it.^37–39^ To investigate whether the observed reduction in OS growth and lung metastasis *in vivo* was mediated by CXCR3A or CXCR3B, we stably transfected the CXCR3 KO mutant with a CXCR3A or CXCR3B expression construct. PCR analysis of the CXCR3 gene revealed transfectant clones with stable transduction of the CXCR3 isoforms (Figure 4A-B). Subsequent CXCL10-mediated chemotaxis assays revealed that CXCR3A-transfected cells were able to rescue the CXCR3 KO phenotype by restoring tumor cell migration to a level similar to that of the parental cell line, whereas CXCR3B-transfected cells failed to rescue the KO phenotype (Figure 4C). Furthermore, utilizing a similar orthotopic xenograft mouse model as described previously, we demonstrated that mice injected with CXCR3A-transfected KO cells developed larger tumors (Figure 4D-E) and significantly more pulmonary metastatic nodules compared to mice injected with CXCR3B-transfected KO cells (Figure 4F), confirming the pivotal function of the CXCR3A isoform in tumor growth and metastasis.

**Figure 4:**
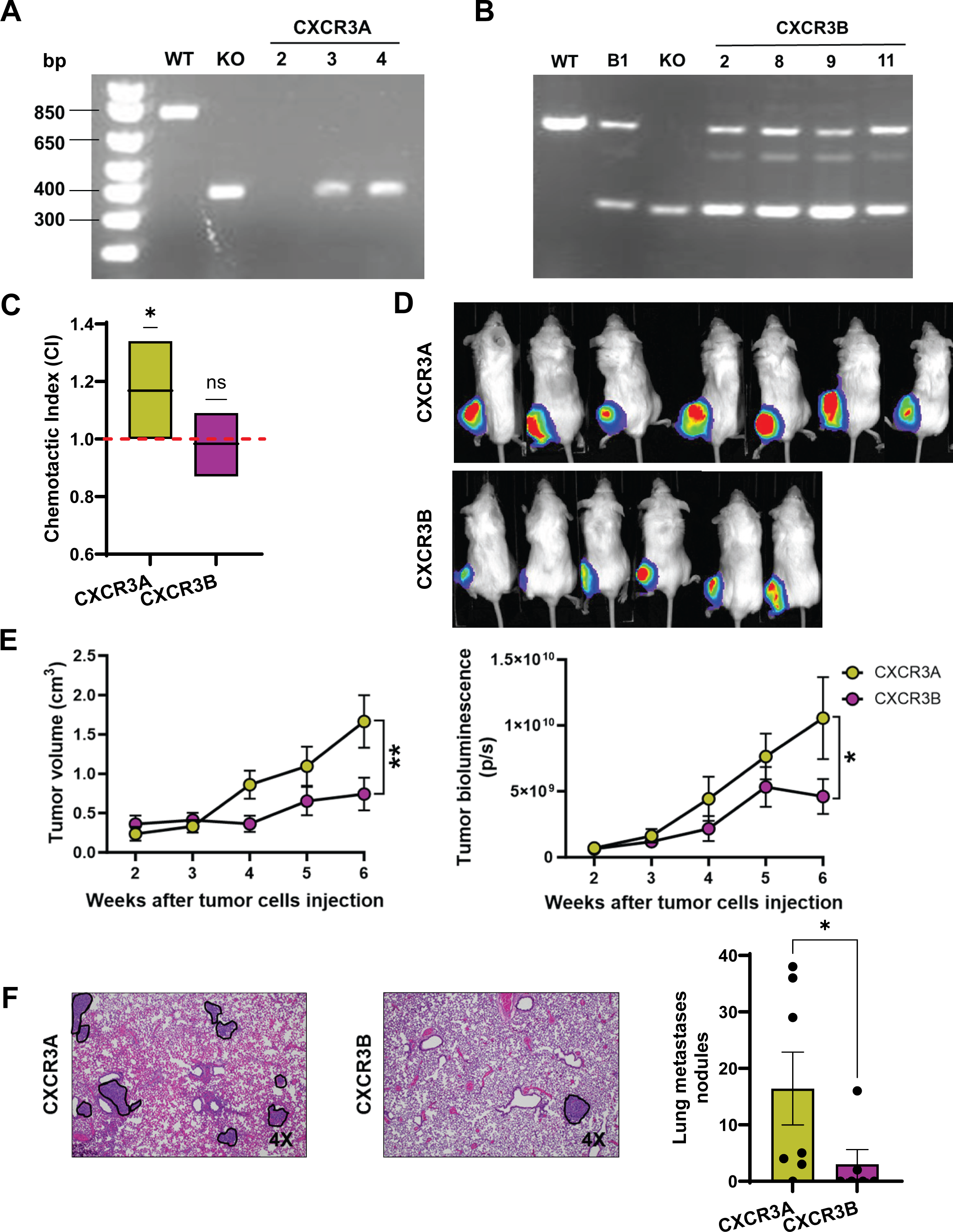
The CXCR3 deletion phenotypes are rescued by the CXCR3A isoform. **A,** PCR analysis of the CXCR3 allele in parental 143B cells, CXCR3 knockout (KO) cells, and CXCR3A transfectants revealed bands of 761 bp, 370 bp, and 370 bp, respectively. Additionally, PCR of the CXCR3 allele in the CXCR3B transfectants produced two bands at 761 bp and 370 bp. For subsequent in vitro and animal experiments, we selected CXCR3A transfectant clone 4 and CXCR3B transfectant clone 11. B1 represents CXCR3B transfectant clone 1. **B,** CXCL10-mediated transwell migration assays of CXCR3A and CXCR3B transfectants from the KO mutant in 0.1% FBS DMEM. **C,** IVIS images of orthotopic xenograft mouse models taken 5 weeks after intratibial injection with CXCR3A (left) and CXCR3B (right) transfectant cells. **D,** Comparisons of primary tumor volumes, calculated as 0.5 x h x w² (left panel), and bioluminescence (right panel) in orthotopic xenograft mouse models of CXCR3A and CXCR3B. **E,** Representative images (left) and quantification (right) of metastatic nodules in H&E-stained resected lungs from the mouse models, with each dot denoting one mouse. Error bars and asterisks represent standard deviations and statistical significance (*P < 0.05; **P < 0.01; ns = not significant).

### Pharmacological inhibition of CXCR3 with AMG487 mitigates aggressive phenotypes in OS

Our findings from the CXCR3 KO mutant suggest that CXCR3 represents a promising therapeutic target for reducing OS metastasis. Given that CXCR3 serves as a common target for all CXCR3 ligands, it presents an attractive option for therapeutic intervention. To assess the therapeutic potential of targeting CXCR3, we utilized a highly potent and selective CXCR3 small molecule antagonist, AMG487.^17,19,20^ We first evaluated the anti-migratory and anti-proliferative effects of AMG487 on three metastatic OS cell lines using transwell migration and CCK-8 proliferation assays, respectively. Our results demonstrated that antagonizing CXCR3 with AMG487 effectively abolished CXCL10-mediated cell migration, reducing it to levels comparable to parental cells in all three OS cell lines (Figure 5A). Notably, AMG487 did not further decrease tumor cell migration in the KO mutant, indicating that the anti-migratory effect of AMG487 is specific to CXCR3 (Figure 5A). This suggests that pharmacologically inhibiting CXCR3 signaling could represent an effective strategy for mitigating OS spread. To validate the inhibitory effect *in vivo*, we employed an orthotopic xenograft mouse model, administering twice-daily treatment of tumor-bearing mice with 5 mg/kg AMG487 for two weeks after tumor injection. Our results revealed that while the treatment had no discernible effect on primary tumor growth (Figure 5B-C), it significantly attenuated the number of lung metastases in mice (Figure 5D). The results indicate the therapeutic utility of CXCR3 targeting in metastatic inhibition for OS.

**Figure 5:**
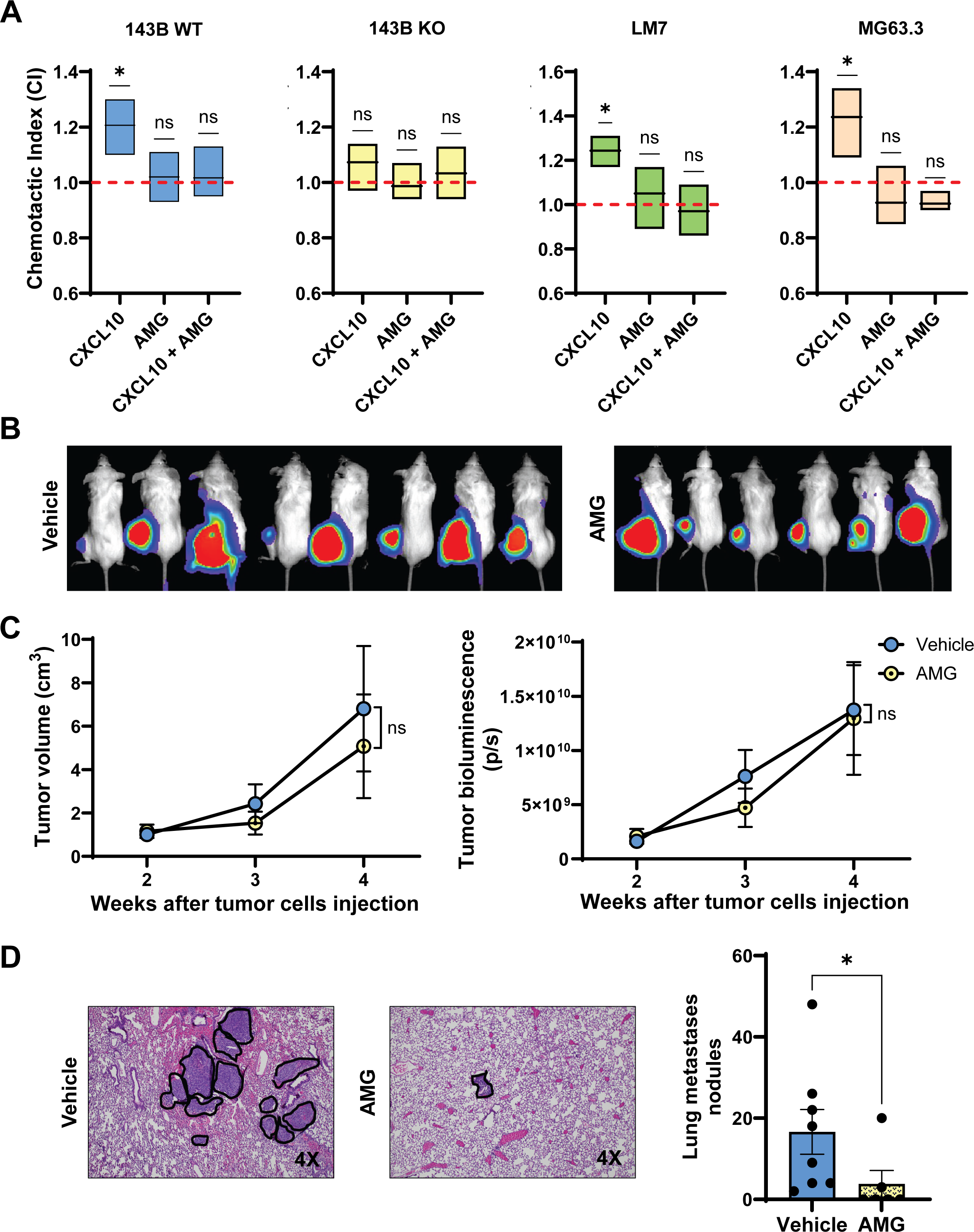
Therapeutic implications of CXCR3 inhibition. **A,** CXCL10-mediated chemotaxis of OS cell lines in transwell migration assays with and without AMG487 (AMG), a potent CXCR3 inhibitor. Overnight starved cells in 0.1% FBS DMEM were treated with 100 ng/ml CXCL10 and 1 µg/ml AMG, with cells pre-incubated with AMG for 1 hour before chemotaxis. **B,** IVIS images of orthotopic xenograft mouse models taken 5 weeks after treatment with vehicle (left panel) and AMG487 (right panel). **C,** Comparisons of primary tumor volumes, calculated as 0.5 x h x w² (left panel), and bioluminescence (right panel) of AMG487-treated or vehicle-treated orthotopic xenograft mouse models of 143B. **D,** Representative images of metastatic nodules in H&E-stained resected lungs from AMG487-treated (AMG) and vehicle-treated mouse models, with each dot denoting one mouse. Error bars and asterisks represent standard deviations and statistical significance (*P < 0.05; **P < 0.01; ns = not significant).

## DISCUSSION

In the current study, we investigated the effects of genetic deletion and therapeutic inhibition of CXCR3 on OS cell proliferation and CXCL10-mediated chemotaxis, as well as the impact of modulating CXCR3 signaling on OS pulmonary metastasis in orthotopic xenograft mouse models. Exogenous CXCL10 increased tumor cell migration in three genetically diverse metastatic OS cell lines with varying levels of CXCR3. CRISPR-mediated CXCR3 deletion in the 143B cell line significantly abrogated the CXCL10-mediated migratory effect, reducing cell proliferation and AKT phosphorylation. Both genetic ablation and pharmacological inhibition of CXCR3 significantly reduced pulmonary metastasis. The CXCR3A isoform rescued the metastatic phenotype in the CXCR3 deletion mutant. Collectively, our findings demonstrate that the CXCL10-CXCR3 axis is crucial for OS metastasis, and targeting CXCR3 can inhibit the development of metastasis.

Survival analysis of CXCR3 and CXCL10 RNA expression data from OS tumor samples in the TARGET dataset reveals that patients with higher expression levels of CXCL10 and CXCR3 have a better prognosis compared to those with lower expression. Since both tumor and immune cells contribute to the bulk tumor used for expression profiling, identifying the source of CXCR3 and CXCL10 expression is crucial. Using a publicly available single-cell RNA sequencing (scRNAseq) dataset, our analysis found that CXCR3 is predominantly expressed in the CD3+ T cell population, while CXCL10 expression is low to undetectable. Therefore, high CXCR3 expression in bulk tumors likely indicates high T cell infiltration, which can enhance the anti-tumor immune response and improve patient survival. Although the scRNAseq data did not identify the exact source of CXCL10 expression, it is evident that it does not originate from tumor cells. Hence, high CXCL10 expression from cells within the tumor microenvironment may elicit paracrine signaling, recruiting a substantial number of CXCR3+ immune cells to bolster the anti-tumor response.

CXCR3 and its cognate ligands—CXCL4, CXCL9, CXCL10, and CXCL11—have been implicated in several malignancies.^16,22,23,37,40–43^ Although these ligands exhibit different affinities for CXCR3 isoforms and elicit distinct functional effects on target cells, our data indicate that all of them produced similar migratory effects on the three OS cell lines tested. These results suggest that targeting CXCR3 is a more effective strategy to inhibit the metastatic behavior of OS cells, as their metastatic potential can be induced by multiple ligands. Additionally, the expression levels of CXCR3 appear to correlate more with the genetic backgrounds of the cell lines rather than their metastatic potential. For example, both MG63 and MG63.3 exhibited low CXCR3 expression, while SaOS-2 and LM7 showed high CXCR3 expression. Conversely, highly metastatic cell lines, such as MG63.3, can have low CXCR3 expression, while less metastatic cell lines, such as HOS, can have high CXCR3 expression. This observation suggests that the metastatic effect of the CXCR3 axis is primarily driven by the availability of its ligands rather than receptor expression. Additionally, the observed phenomenon may be affected by the cell surface expression of CXCR3 rather than total cellular protein levels.

While our findings suggest that the CXCR3 signaling pathway is crucial for OS pulmonary metastasis, recent reports have also highlighted the involvement and cross-talk of the CXCR4 and CXCR7 axes in other tumor metastases.^44–46^ The upregulation of the CXCL12-CXCR4 axis has been found to increase metastases in the lungs for melanoma^47^ and in both lymph nodes and lungs for breast cancer.^48^ Additionally, chronic lymphocytic leukemia (CLL) patients with a CXCR3^low^/CXCR4^high^ phenotype exhibited more aggressive disease and poorer outcomes.^49^ In colorectal cancer (CRC) cells, CXCR3 expression prevented CXCR4 internalization, thereby promoting CRC invasiveness, suggesting potential cooperativity between these receptors.^18^ In OS, CXCR4 expression has been shown to promote proliferation and lung metastasis by downregulating miR-613 and inhibiting apoptosis.^50,51^ Interestingly, CXCR3 and CXCR4 can form functional dimers, although this feature has yet to be characterized in OS.^52^

Strategies to target metastasis via CXCR3 antagonism have garnered significant attention recently.^17,19,20^ Using AMG487, a highly specific and potent CXCR3 antagonist, we demonstrated that systemic administration of AMG487 can significantly impede the development of lung metastasis originating from primary tumors located in the tibia in an orthotopic xenograft mouse model. This model effectively captures the full metastatic cascade process, which is a major improvement from the previous study using an experimental metastasis model focused solely on lung colonization.^17^ Other studies have also reported similar anti-metastatic effects of CXCR3 antagonism in various solid malignancies, further supporting the clinical potential of CXCR3 antagonism in the anti-metastatic treatment of OS.^19,20^ Another novel finding in our study was that CXCR3 deletion, unlike AMG487 treatment, reduced OS cell proliferation *in vitro* and markedly impeded primary tumor growth in mice. This suggests that CXCR3 signaling is crucial for tumor growth. The differential effects between the deletion mutant and the antagonist indicate that CXCR3 signaling may be vital during the early stages of tumor growth or establishment but its influence diminishes once the tumor is already established.

Given that host immune cells, such as activated CD8+ T cells and NK cells, can be trafficked to tumor sites via CXCR3,^13–15^ one potential concern with using a CXCR3 antagonist is the possibility of creating an immune-depleted or desert tumor microenvironment in OS. However, using immunocompetent mouse models, Pradelli *et al*. demonstrated that control mice developed more lung metastases compared to AMG487-treated mice, indicating that CXCR3 antagonism provided better metastatic control in the presence of a fully functional immune system.^17^ Similarly, Walser *et al*. observed in a murine model of metastatic breast cancer that the anti-metastatic effects of AMG487 were dependent on NK cells.^19^ Since NK cells express CXCR3,^53^ the report argued that the anti-metastatic effects could involve CXCR3-null NK cells, among other factors. Contrary to the belief that CXCR3 is essential for T cell tumor infiltration, Chow *et al*. found no difference in the absolute numbers of effector CD8+ T lymphocytes between CXCR3-/- melanoma mice and controls, suggesting that CXCR3 primarily influences T cell positioning with CXCL10-expressing dendritic, macrophage, or Th1 cells, rather than controlling intratumoral infiltration.^54,55^ These findings indicate that the immunomodulatory effects of CXCR3 blockers do not undermine their anti-metastatic function. Nonetheless, further studies are necessary to elucidate the precise interplay between CXCR3 antagonism and intratumoral immune cell functions, including those of macrophages, T cells, and NK cells. Moreover, fewer than 25% of OS cases are classified as immune-hot tumors.^56^ The use of CXCR3 antagonists in immune-cold OS may therefore produce a strong anti-metastatic effect while minimizing potential negative impacts on immune response.

An advantageous aspect of CXCR3 inhibition lies in the recent development and utilization of a clinically applicable antagonist. ACT-777911, a potent, unsurmountable CXCR3 competitive inhibitor, boasts well-established efficacy and safety profiles across multiple species, including mice, rats, dogs, monkeys, and humans.^57,58^ Its successful First-in-Human adult clinical trial demonstrated excellent properties in terms of absorption, distribution, metabolism, and excretion (ADME), as well as efficacy and safety.^59^ While initially developed for treating recent-onset type 1 diabetes, our study’s findings support its potential application in OS metastasis treatment. Currently, our laboratory is assessing the anti-metastatic effects of ACT-777911 through various *in vitro* and animal models of OS, including patient-derived xenograft models.

Drawing on our current and prior investigations, alongside relevant studies from other researchers, we propose a working model to elucidate the functions of the CXCL10-CXCR3 axis in OS (Figure 6). Elevated levels of CXCL10 within the tumor microenvironment stimulate OS cell migration while simultaneously recruiting CXCR3+ immune cells to the tumor site. Subsequently, heightened OS cell migration facilitates metastatic colonization of distant organs, particularly the lungs, leading to tissue damage and inflammation, thereby further elevating circulating CXCL10 levels. In a recent study, Pein *et al*. observed that metastatic breast cancer cells can prompt lung fibroblasts to upregulate CXCL9 and CXCL10 expression, fostering an inflammatory microenvironment via the NF-kB pathway to promote lung metastases. ^60^ Disruption of the tumoral CXCL9/CXCL10-CXCR3 axis notably hindered metastatic colonization. Additionally, our recent findings indicated that increased circulating CXCL10 concentrations correlated with poorer outcomes in OS patients, suggesting that the chemokine might exert a chemoattractant effect on anti-tumor immune cells, diverting them away from the tumor site and potentially enhancing the ability of OS cells to evade immune clearance. This hypothesis is supported by our data that elevating circulating CXCL10 levels in immunodeficient orthotopic xenograft mouse models of OS did not lead to increased pulmonary metastases (Supplementary Figure S3), implying that the circulating chemokine might not directly influence tumor cells. However, further investigations utilizing immunocompetent mouse models are warranted to dissect the role of circulating CXCL10 in OS and immune cells.

**Figure 6:**
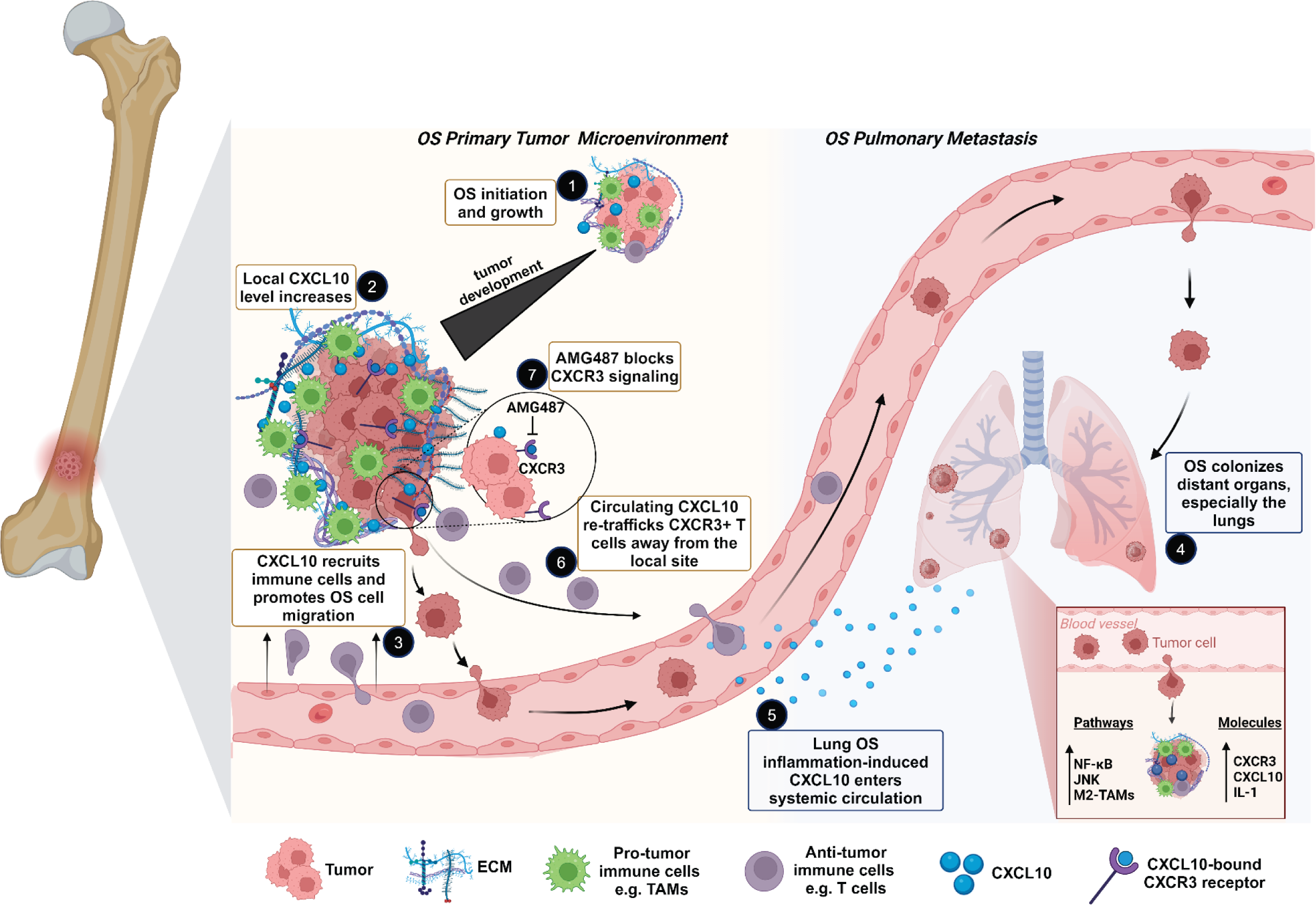
Working model of the CXCL10-CXCR3 axis in OS. Elevated levels of CXCL10, produced by tumor cells or other unknown sources within the tumor microenvironment, play a crucial role in OS progression. CXCL10 promotes OS cell migration and attracts CXCR3+ immune cells to the tumor site. Consequently, increased tumor cell migration leads to metastatic colonization of the lungs. This process is facilitated, in part, by the polarization of M2-like tumor-associated macrophages (TAMs) and IL-1 expression via JNK signaling. The resulting NF-κB-mediated inflammatory phenotype in lung fibroblasts causes tissue damage and further amplifies the production of circulating CXCL10. Higher circulating CXCL10 levels serve as an indicator of metastasis, recruit CXCR3+ immune cells away from the tumor site, and correlate with worse outcomes in patients. Notably, our model and data demonstrate that blocking CXCR3 using AMG487 effectively inhibits metastasis in OS. The figure was created with BioRender.com.

In summary, we have provided strong evidence that highlights the critical role of CXCR3-CXCL10 signaling in the progression of pulmonary metastases in OS using orthotopic xenograft mouse models. Our findings from both CXCR3 deletion mutant and AMG487 treatment experiments underscore the therapeutic potential of targeting CXCR3 signaling in inhibiting OS metastasis. Further investigations into CXCR3 inhibition, employing clinically applicable antagonists and patient-derived xenografts, hold promise for the development of novel and effective therapeutic interventions for metastatic OS, potentially ameliorating the current poor prognosis associated with this condition.

## AUTHOR CONTRIBUTIONS

BG, JW, XC, MW, ZDM, and TKM performed experiments and/or analyzed data. AM and JH conducted the histological analysis of mouse samples. MW and JX generated the CXCR3 KO mutant. TKM conceived the project, designed the study, and supervised the experiments. BG and TKM drafted the manuscript. All authors reviewed and revised the manuscript and approved its content.

## ACKNOWLEDGMENTS

We would like to thank Weidong Jin for his technical assistance with the mouse study. We also acknowledge the TARGET OS consortium for generating bulk RNA expression data. Generous funding support was provided by the Kate Amato Foundation and the Cancer Prevention and Research Institute of Texas (RP200135) to TKM.

## Supplementary Figures

**Supplementary Figure S1:**
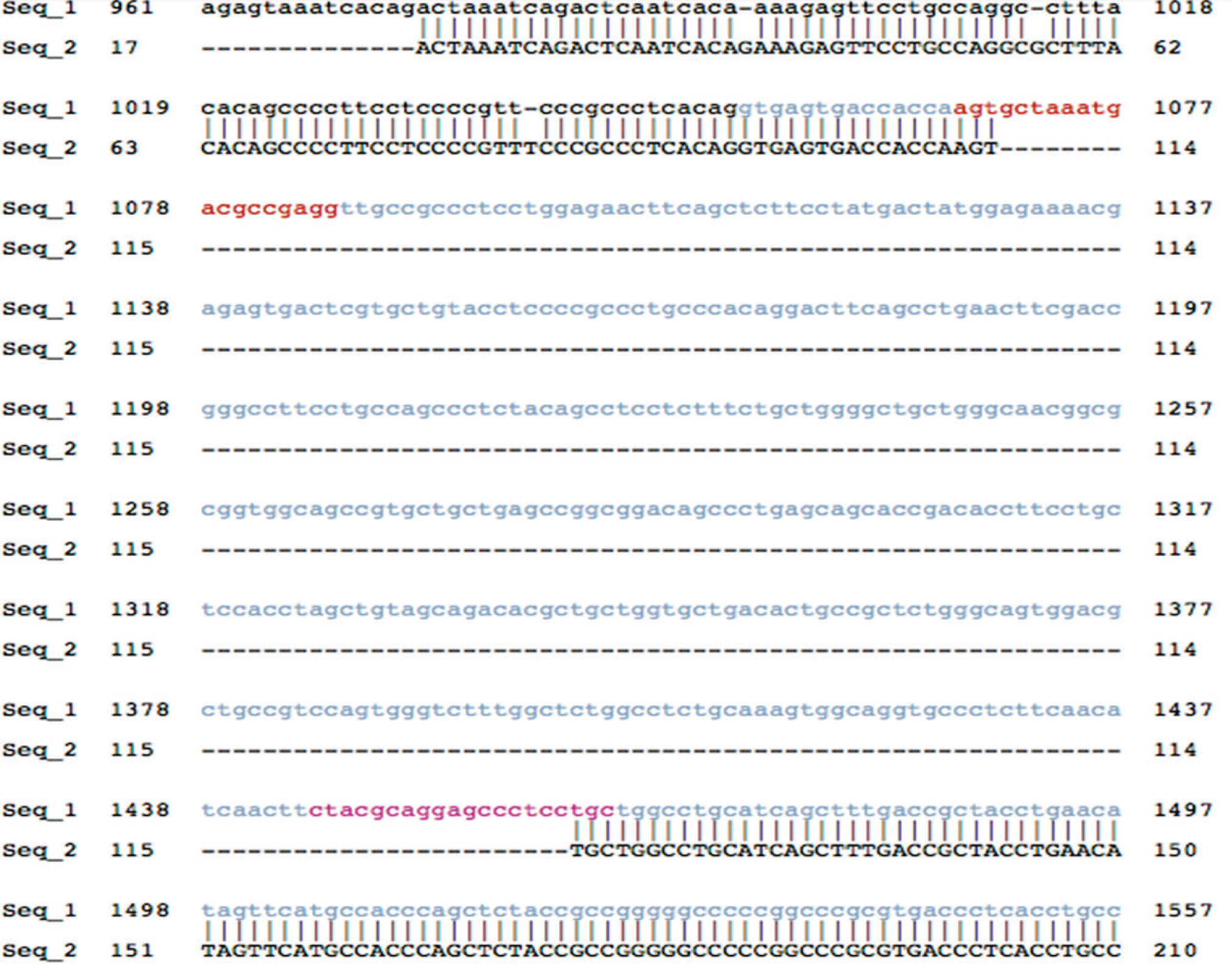
Sequence alignment reveals a CXCR3 deletion in 143B cells. The top sequence (seq_1) corresponds to the reference genomic sequence of CXCR3, while the bottom sequence (seq_2) represents the mutant CXCR3 sequence in the CXCR3 knock-out mutant.

**Supplementary Figure S2:**
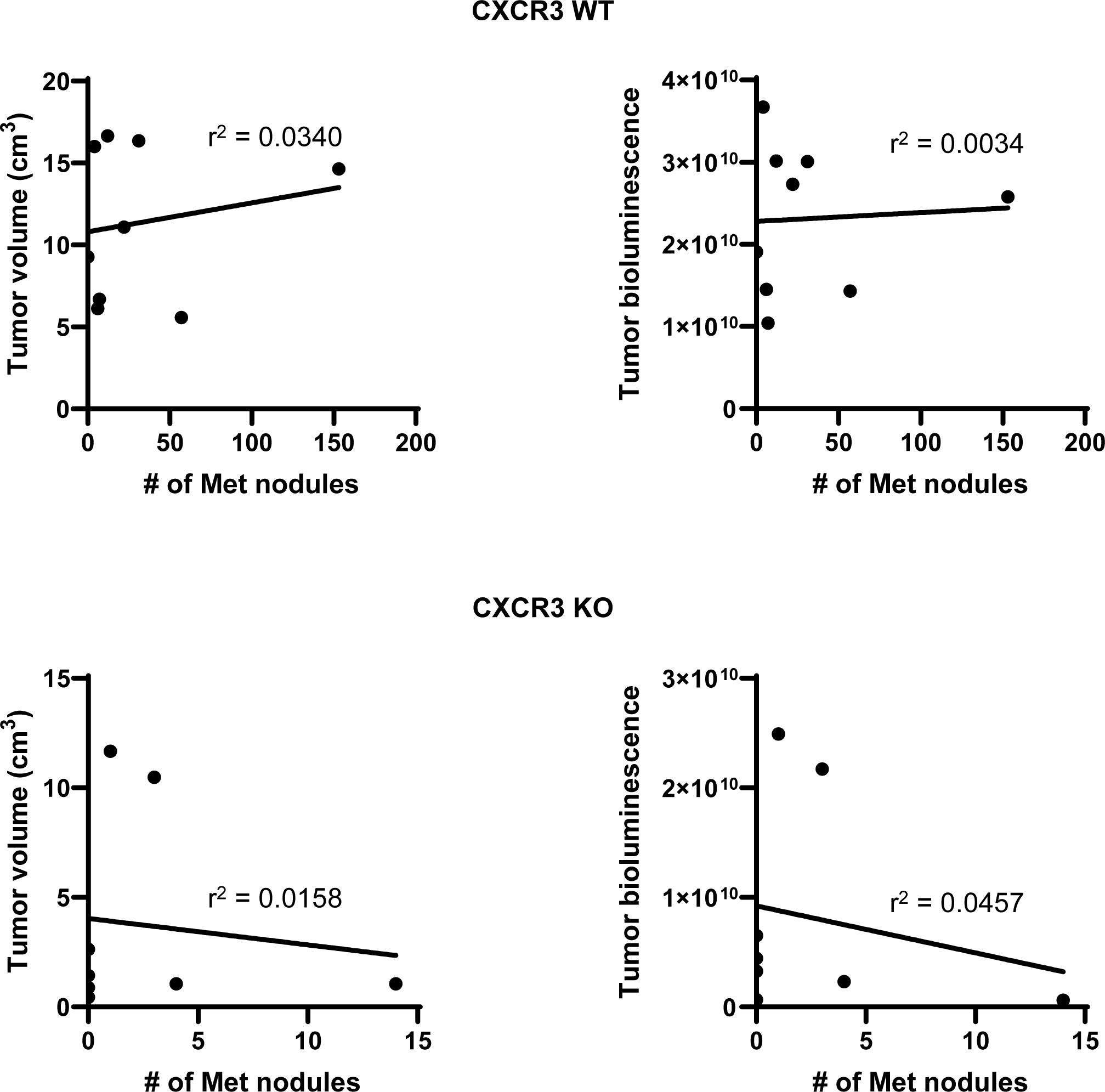
Correlation between lung metastasis rate and primary tumor volume or activity in orthotopic xenograft mouse models. Mice were intra-tibially injected with either 143B parental cells or CXCR3 knockout (KO) cells. Over a 5-week period, we monitored tumor growth using IVIS (*in vivo* imaging system). Primary tumor volumes were calculated as 0.5 × h × w² (where h represents height and w represents width). Additionally, we quantified the number of metastatic nodules in resected lungs from the mouse models using H&E staining. Each data point corresponds to one mouse. Pearson correlation coefficients were computed to assess the relationship between lung metastasis rate and primary tumor characteristics.

**Supplementary Figure S3:**
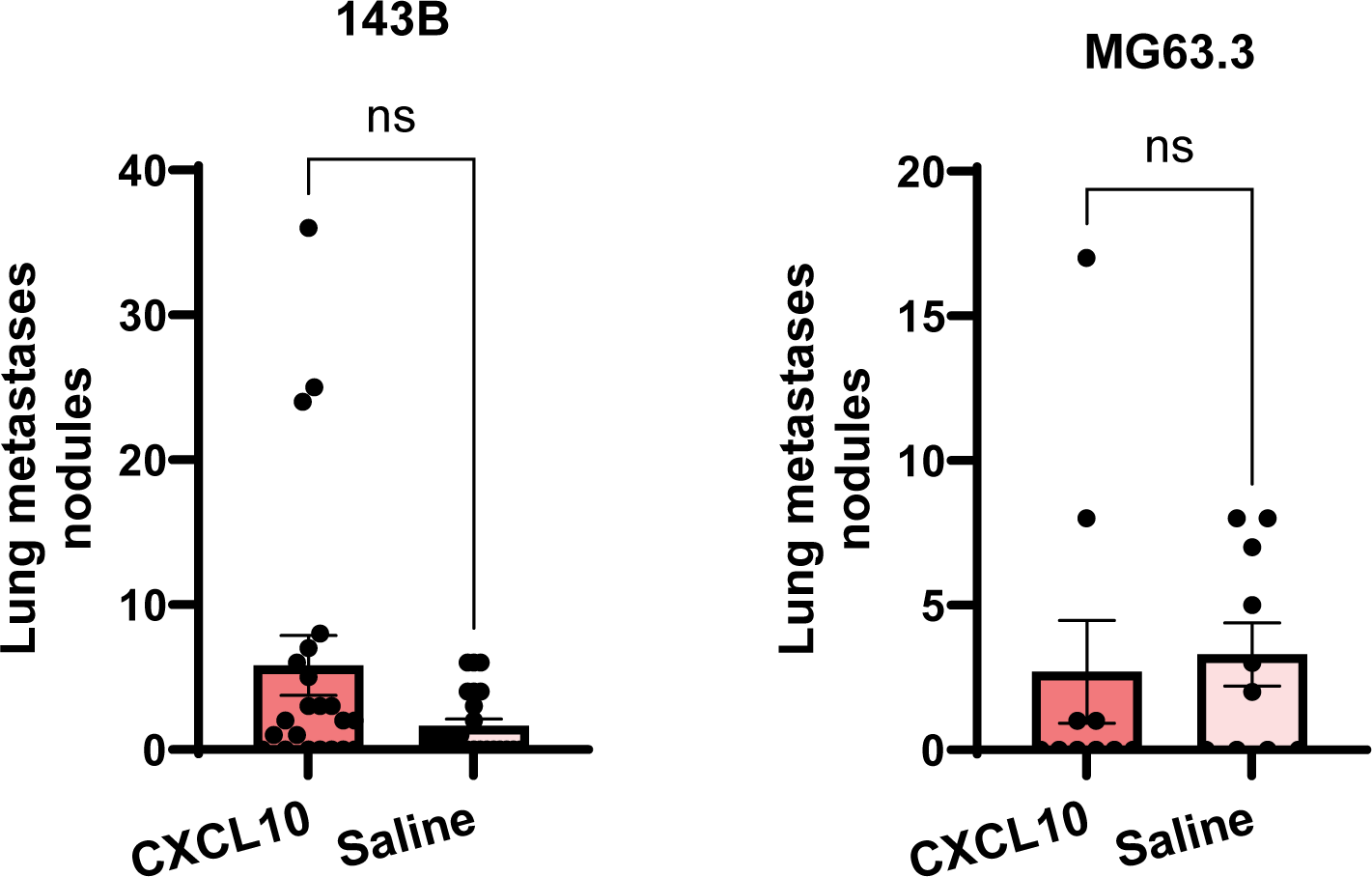
Orthotopic xenograft mouse models investigating circulating CXCL10 effects on osteosarcoma metastasis. We utilized orthotopic xenograft mouse models to explore the impact of circulating CXCL10 on osteosarcoma metastasis. The scatter dot plot illustrates the number of metastatic nodules identified in mouse lungs resected from the CXCL10-treated group and the control group. Mice were tibially injected with either 143B or MG63.3 cells and treated daily with 1.5 µg of CXCL10 or saline intravenously for two weeks. The study endpoint was the detection of luminescence in the lungs of control mice. Two batches of experiments were conducted for 143B cells and combined, while one batch was performed for MG63.3 cells. Statistical differences were assessed using the two-tailed Mann-Whitney test. Error bars and asterisks denote standard deviations and statistical significance, respectively (ns = not significant)

